# Models of microbiome evolution incorporating environmental microbial selection

**DOI:** 10.1101/2024.10.17.618834

**Authors:** Yao Xiao, Teng Li, Allen Rodrigo

## Abstract

The environment plays a crucial role in shaping the microbial composition of hosts. Different environments harbor distinct microbial communities, leading to variation in the availability of different microbes to potential hosts. Over the past few decades, human activities have significantly altered the environment, resulting in a decline in microbial diversity. Currently, the impact of this environmentally-induced reduction in microbial diversity on environmental microbial communities, host-microbiome interactions, and their evolutionary implications remains unclear. In this paper, we simulated how environmental microbial selection can lead to decreasing microbial diversity, using a previously-developed selection model. Our results indicate that microbial communities are influenced by both environmental conditions and host preferences. The diversity of the microbiome carried by hosts is impacted by the environment, with stronger environmental microbial selection reducing microbial diversity in the hosts, while at the same time, facilitating the emergence of mutually beneficial symbioses between microorganisms and hosts. Environmental microbial selection (EMS) leads to the survival of highly fit microorganisms in the environment, while the fitnesses of hosts and microbes within hosts remain relatively unaffected by environmental microbial selection.

## Introduction

The composition of microbial communities in the environment is an important factor influencing the microbiomes of hosts. Structural variation in the environment, due to graded or step differences in biotic and abiotic properties, can lead to differences in the environmental microbial content [1]. For instance, the bacterial communities in the atmosphere vary between agricultural fields, suburban areas, and forests [2]. Similarly, the composition of soil microbial communities differs across different types of forests [3]. This phenomenon is also observed in aquatic environments, where microbial communities in the branches and main streams of rivers within the same region exhibit significant differences [4]. Even in urban areas, specific taxonomic groups of microorganisms in the air differ between cities [5].

The composition of microbiomes in the host is expected to vary depending on the environment in which the host lives. Some researchers have shown that the environment has a more significant role in shaping the human gut microbiome than host genetics [6]. Broadly, the diet of the host is also considered a part of the environmental composition and can influence the composition of the host’s microbiome [7, 8]. Rural children from Burkina Faso, who have a diet rich in fiber compared to European children, exhibit significant differences in their gut microbial composition [9]. Smoking and alcohol consumption also influences the human microbiome. Compared to non-smokers, smokers have lower diversity in their oral and gut microbiomes [10, 11]. Heavy alcohol drinkers, on the other hand, show an increased abundance of bacteria from the phylum Proteobacteria and a decreased relative abundance of bacteria from the phylum Bacteroidetes in their colon microbiome [12]. However, red wine polyphenols have been found to increase the abundance of bacteria from the phyla Proteobacteria, Firmicutes, *Actinobacteria*, and *Bacteroidetes*, while consuming pine nut wine has the opposite effect [13].

Environmental influences on the host microbiome affect non-human animals as well. It has been shown that air pollutants affect the microbiome of the intestinal tract of mice [14]. During the winter migration of swan geese from the north to the south of China, the gut microbiome changes when they arrive in different areas [15]. Differences in the level of contact with humans can also lead to changes in the composition of microbiomes in animal hosts. Microbiome components that are abundant in humans are present in captive mammals but not in the microbiome of individuals of the same species in the wild [16–18]. Another study found that by comparing the gut microbiomes of coyotes (*Canis latrans*), crested anole lizards (*Anolis cristatellus*), and white-crowned sparrows (*Zonotrichia leucophrys*) in urban and rural populations, researchers were able to find human gut microbial components in the gut microbiomes of urban animal populations [19]. Thus, the environment is an essential factor influencing the microbiome in the host.

In recent years, various studies have shown a decrease in microbial diversity in the environment due to frequent human activities. For example, urbanization has been shown to result in lower bacterial diversity in urban air than in rural areas [20]; increased land use has resulted in decreased microbial diversity in alpine temperate forest soils [21]; and climate change and long-term warming can lead to a decrease in soil microbial diversity in grasslands [22], and increased drought levels can also lead to a decrease in soil microbial diversity [23]. It is not clear exactly what effect reduced diversity in the environment will have on long-term host and microbial coevolution.

In our previous research [24], we extended Zeng et al’s selection model [25] by incorporating a resource provisioning process to investigate how much fitness the host is willing to invest in adjusting its own microbiome as it coevolves with microbes. Our results indicate, in part, that the composition of the environmental microbiome is an essential factor influencing the extent to which the host sacrifices resources. In this paper, we include environmental microbial selection (EMS) in the Zeng et al. model [25] to explore the effect of the microbial selection by the environment on long-term microbial and host coevolution.

## Model

In Zeng et al’s selection model [25], each microbial species/Operational Taxonomic Unit (OTU) is assigned a certain number of traits in a phenome, and each trait can affect the fitness of OTUs and hosts independently. So each trait is assigned two independent values; one represents the effect of the trait on the fitness of hosts, and the other represents the effect of the trait on the fitness of OTUs. The values that these traits can take are -1, 0, or 1, where -1 indicates that the trait has a negative impact on fitness (of hosts or OTUs), 0 indicates that the trait has no effect on fitness and is therefore neutral, and 1 indicates that the trait has a positive effect on fitness. Based on these traits, the two fitness contributions of each OTU can be calculated. These two contributions influence the degree of host selection (HS) and microbial selection (MS), respectively. Under HS, hosts with microbes that collectively possess more positive-valued host-directed traits have a higher probability to survive or reproduce relative to hosts with microbes that have fewer positive-valued traits. Similarly, under MS, OTUs with more positivevalued microbe-directed traits have a higher probability of surviving or reproducing when compared to OTUs that possess fewer such traits.

Our extension [24] to Zeng et al [25] included a modifier, *k*, representing the amount of fitness the host sacrifices to enhance or reduce the fitnesses of microbes that contribute positively or negatively to the host, respectively. Readers are directed to [24] for the details on our model incorporating this Resource Provisioning Process (RPP), and how *k* is used in the fitness calculations.

In the model we develop in this paper, we use Yao et al’s RPP model, and we modify each OTU’s phenome by assigning a third independent value to each trait to accommodate the effect of the trait on the environmental microbial fitness of OTUs. These traits can again take values of -1, 0, or 1, where -1 indicates that the trait has a negative impact on the environmental fitness of OTUs, 0 indicates that the trait has no effect on the environmental fitness of OTUs and is therefore neutral, and 1 indicates that the trait has a positive effect on environmental fitness of OTUs. Based on the traits, we can calculate the contribution to the environmental fitness of each OTU. Microorganisms with higher environmental microbial fitness are more likely to survive and reproduce in the environment. Also, in our model, the composition of the environment is composed of microorganisms derived from hosts (y%) and the fixed environment (1-y%). Once our model calculates this mixed environment, EMS is performed. The frequency of OTUs in the final environment is calculated based on the environmental microbial fitness of different OTUs and the frequency of OTUs in the mixed environment.

An example is shown here to illustrate EMS (Fig. 1). In this example, we have three hosts, three microbial species, and three traits available to microbes. For illustrative purposes, each microorganism’s phenome has two traits. Based on the microbiome of the host, we can calculate the pooled contribution from hosts to the environment (= “the pooled environment”). The mixed environment consists of 50% of the pooled environment and 50% of the fixed environment. Finally, we can get the final environment after EMS. We can find that after the incorporation of EMS, the frequency of microorganisms with high environmental microbial fitness is significantly increased, and the frequency of microorganisms with low environmental microbial fitness is significantly decreased.

**Figure 1:**
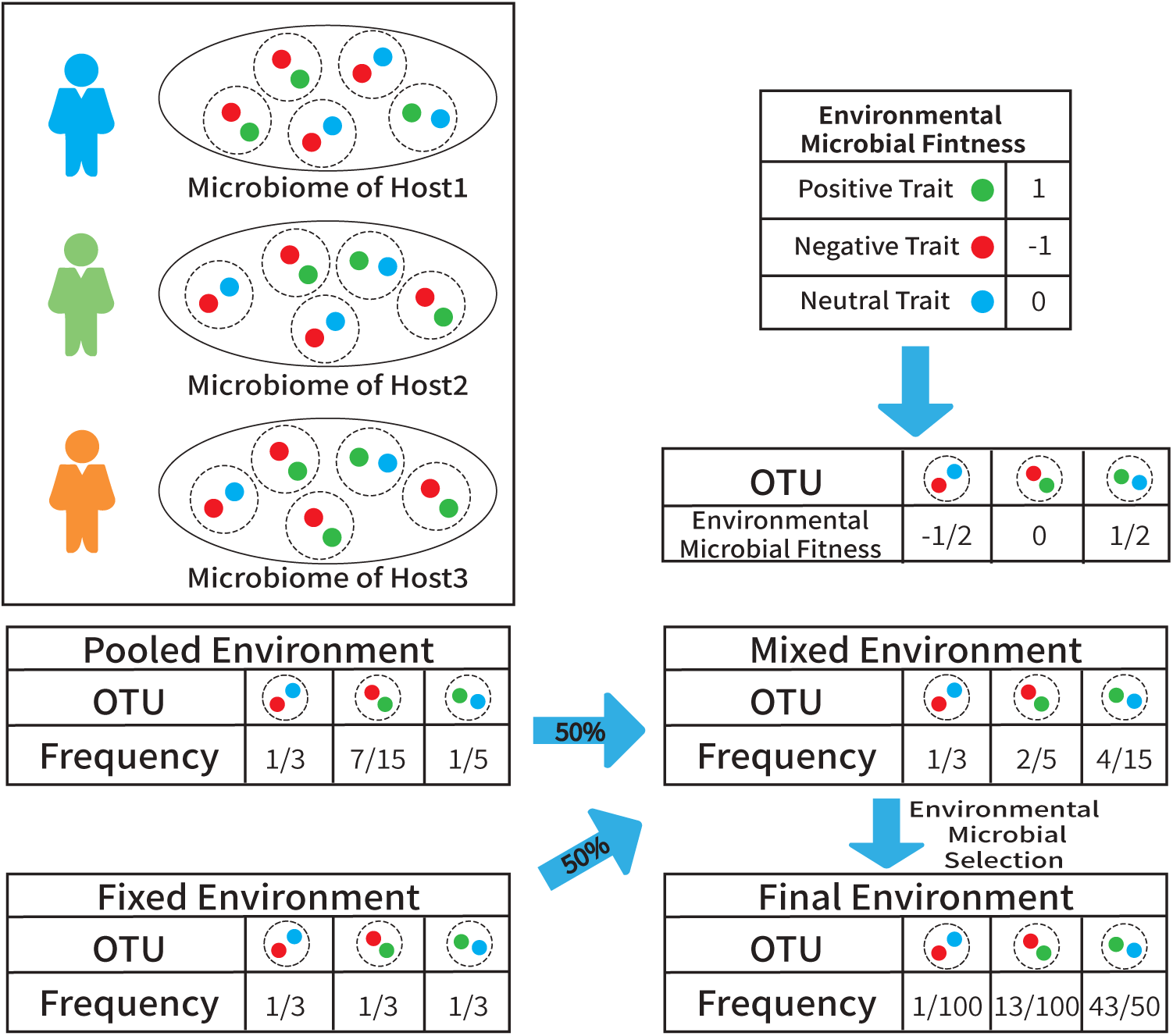
An example of EMS. This figure shows the change in the microbial composition of the environment with the incorporation of EMS. In this example, there are three microorganisms, each given two characteristics. The color of the feature represents its effect on the survival of the microorganism in the environment (green is positive, red is negative, and blue is neutral). Based on the characteristics of each microorganism, we can calculate its environmental microbial fitness. Meanwhile, the mixed environment here consists of 50% pooled environment and 50% fixed environment. We can calculate the pooled environment based on all the microorganisms carried by the host in the host population. Finally, the final environment is calculated by EMS based on the environmental microbial fitness and mixed environment.

## Results

When we compare the simulation results without resource provisioning (RPP), we find that the differences in microbial diversity, microbial composition, and fitness between the simulations without RPP and with RPP are negligible. Therefore, we present only the simulation results with RPP. Also, we applied different intensities of microbial selection, host selection, and environmental microbial selection for the simulations. In our previous research, changes in the intensity of HS do not cause substantially large changes in diversity. For this reason, we only report the results when *S_HS_* = 10. The simulation results without RPP and other results with different HS intensities are available in *Supplementary Information*.

### Microbial Diversity in Environment

First, our model is able to record *α*-diversity of the microbial community in the environment. We find that in the absence of EMS (*S_EMS_* = 1), the addition of MS (e.g., *S_MS_*= 10, 100, 1000) decreases *α*-diversity in the environment, especially when the percentage of host contribution to a pooled environment (PE) is high (Fig. 2). This is because when PE is high (i.e., *PE* = 90, 100), the environmental microbiome is highly influenced by the host microbiome. At the same time (when *PE* = 90, 100), the microbial diversity in the host is reduced due to MS (Fig. 3). So when PE is high, microbial diversity in the environment decreases as microbial diversity in the host decreases.

**Figure 2:**
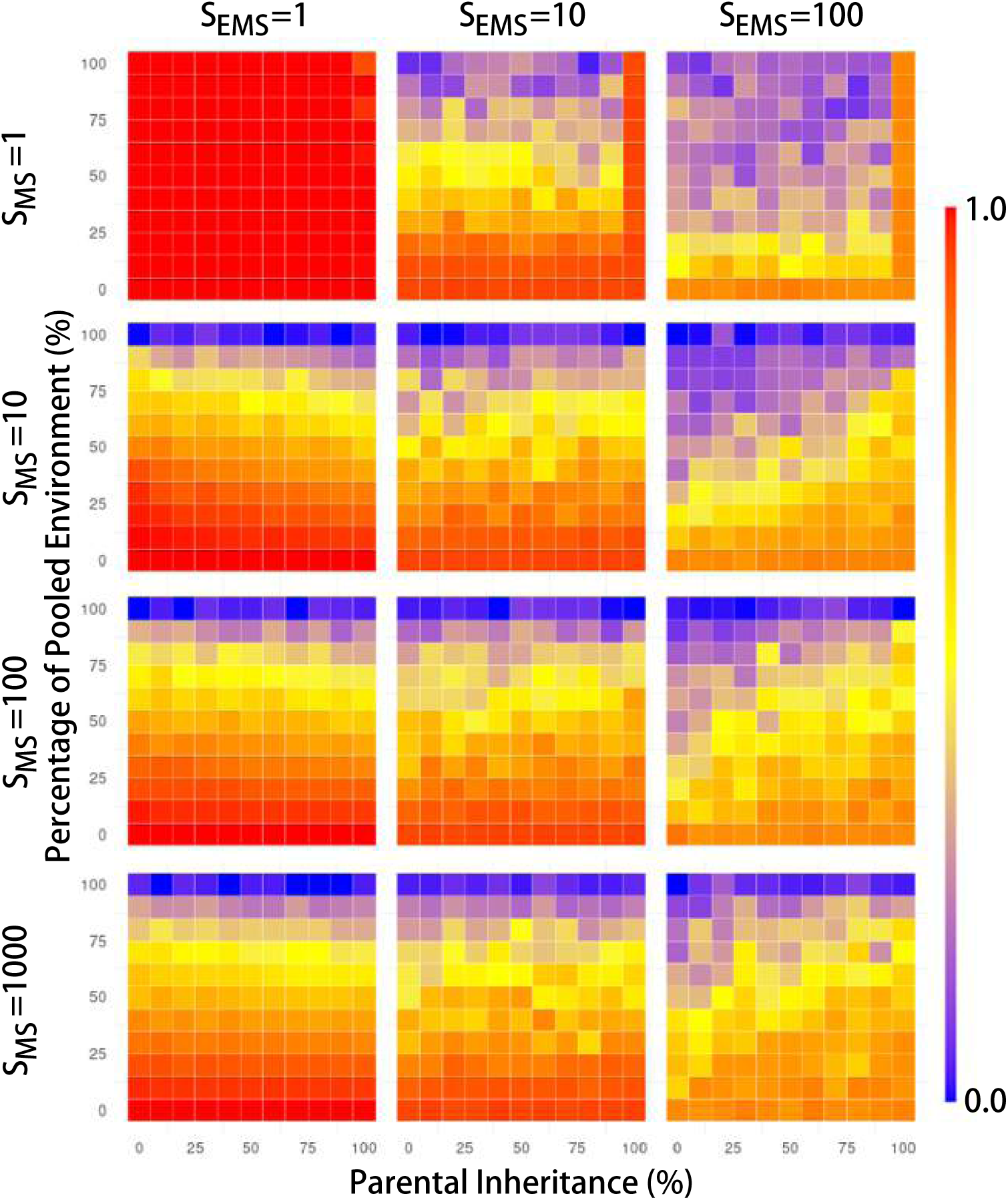
Pattern of *α*-diversity in the environment (*S_HS_* = 10). Each heat map represents its corresponding combination of EMS and MS. There are three levels of EMS: 1, 10, 100, and four levels of MS: 1, 10, 100, 1000. for each heat map, the horizontal axis is the parental contribution to offspring (i.e., the parental inheritance, PI), and the vertical axis is the parental contribution to the environment (i.e., the percentage of pooled environment, PE). The scale is linear, from 0 to 100. The color bars on the right side represent the corresponding diversity values (blue for low diversity, yellow for medium diversity, and red for high diversity).

**Figure 3:**
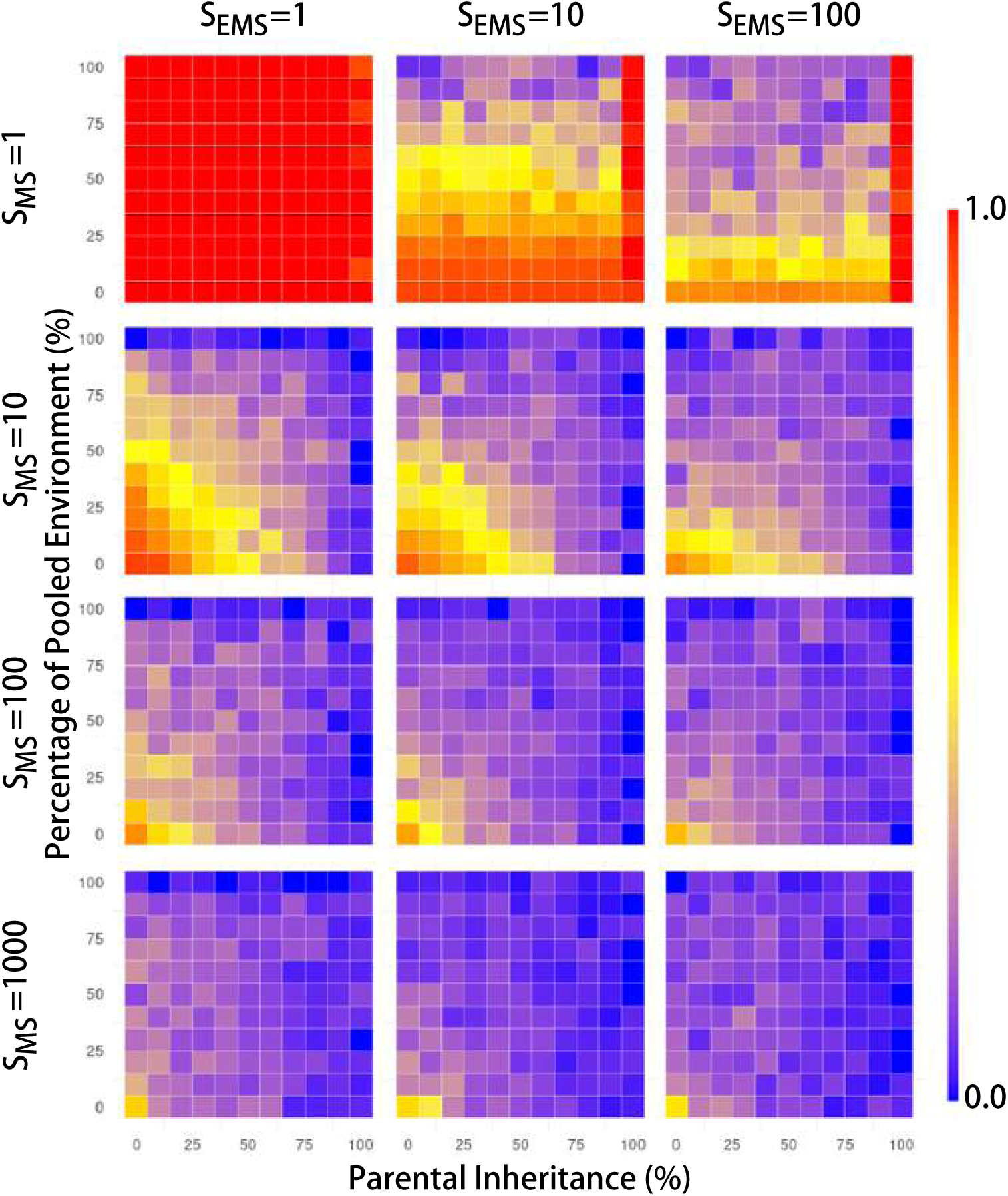
Pattern of *α*-diversity in the hosts (*S_HS_*= 10). This figure is plotted in the same way as Fig. 2.

When introducing EMS (*S_EMS_* = 10, 100), it is easy to see that *α*-diversity decreases as the intensity of EMS increases (Fig. 2). This is reasonable because strengthening EMS will more easily lead to the disappearance of some microorganisms that are less adapted to the environment. At the same time, for each combination of *S_MS_* and *S_EMS_*, *α*-diversity decreases as PE increases (Fig. 2). This is because when PE is low, the source of microorganisms in the environment is mainly from the fixed environment; in contrast, when PE is high, microbes are principally contributed by the host.

Surprisingly, in the case when *S_EMS_ >* 1, and *S_MS_ >* 1, *α*-diversity tends to increase with increasing intensity of MS when PE and the parental inheritance (PI) are high. We hypothesize that the combination of high MS, PE and PI leads to a significant contribution by the hosts of a subset of microorganisms that are strongly selected in the hosts, to the environment. Therefore, when the host microbiome is able to influence the environmental microbiome, some microorganisms that are not dominant in the environment but are dominant in the hosts can also survive in the environment. As a result, *α*-diversity in the environment is increased.

Finally, it’s interesting to note the pattern of *α*-diversity in the environment when there is no MS (*S_MS_*= 1) and PI is 100% (*PI* = 100). Since there is no MS and the host does not acquire microorganisms from the environment, the host microbiome is unaffected by MS and EMS. So *α*-diversity in the host is always high (*S_MS_*= 1 and *PI* = 100), and hardly any microorganisms ever go extinct in the host – these results are also seen in Figures 3 and 4 of [25]. If the environment acquires microorganisms from the host, then the host microbiome can keep replenishing microorganisms so that the environmental microbial diversity is always high.

**Figure 4:**
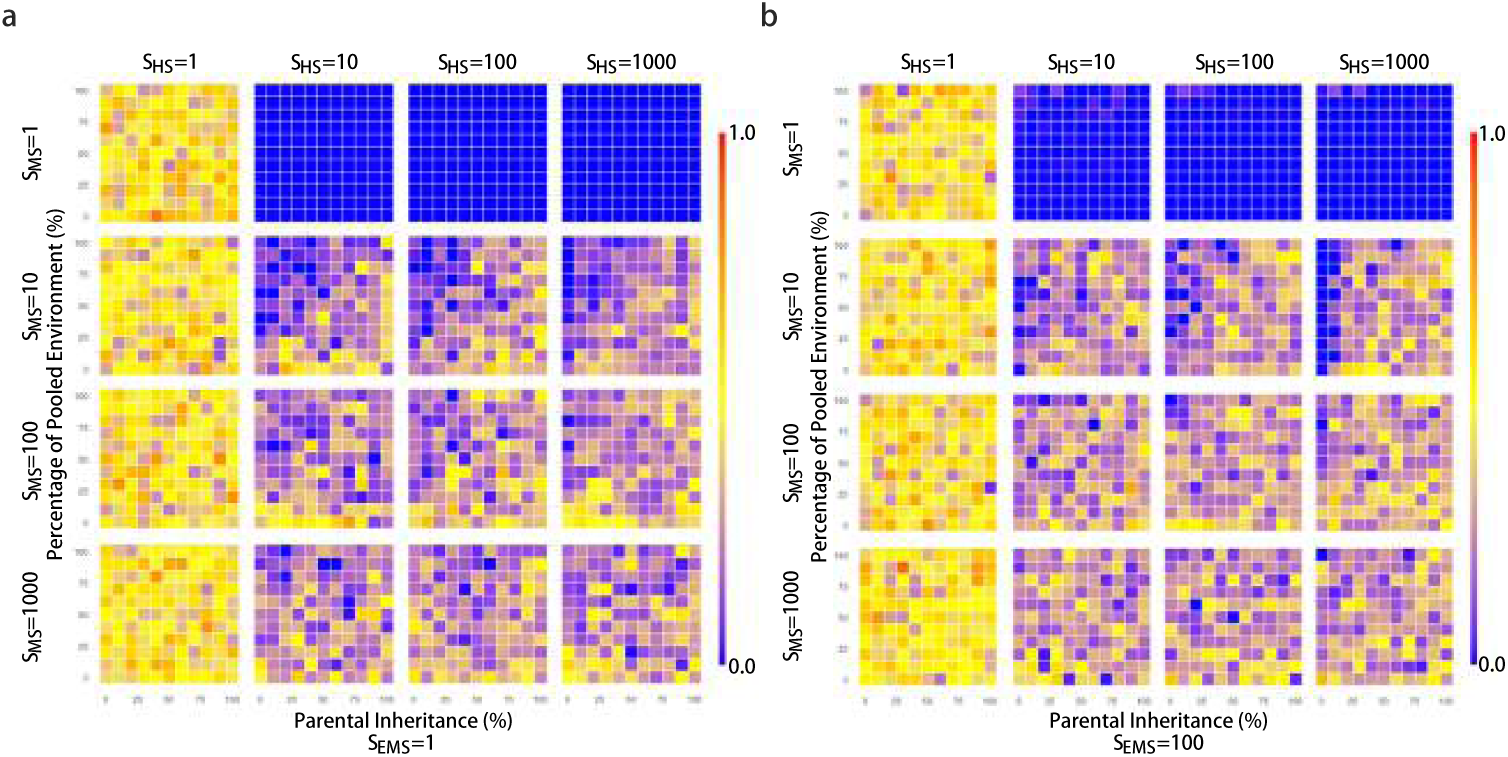
Pattern of *k* without (Fig. a) and with (Fig. b) EMS. Each figure is plotted in the same way as Fig. 2, but the color represents the corresponding the values of *k*. In addition, Fig. a is when *S_EMS_* = 1 (without EMS), and Fig. b is when *S_EMS_* = 100 (with EMS).

### Microbial Diversity in Hosts

Next, we analyzed the diversity of microorganisms within the host (Fig. 3). We focus on *α*-diversity because *β*-diversity is always low (Supplementary Information: Figure S9), while the heatmaps of *α*-diversity and *γ*-diversity are almost identical (Supplementary Information: Figure S10 and Figure S11), consistent with the findings of Zeng et al. (2017) [25].

Interestingly, *α*-diversity decreases as both intensities of MS and EMS increase (Fig. 3). In the absence of EMS (i.e., when *S_EMS_*_=1_), the results obtained are similar to those obtained by Zeng et al. (2017) [25]: as *S_MS_* increases we see a reduction in *α*-diversity, because microorganisms are under increasing intensities of selection. When we introduce EMS (*S_EMS_ >* 1), *α*-diversity further decreases as the intensity of EMS increases, especially when PI is low (i.e., *PI* = 0, 10, 20). This is because when PI is low, the host acquires most of the microorganisms from the environment. So the microbiome in the host is highly influenced by the environment.

The patterns of *α*-diversity of the microbiome in environment and hosts are identical when there is no MS (*S_MS_* = 1; compare Fig. 2 with Fig. 3 above). This is reasonable because the absence of MS does not filter out some microbes. However, When hosts acquire all the microorganisms from the parent (*PI* = 100) and EMS is present (*S_EMS_ >* 1), the diversity of the microbiome in hosts is significantly higher than that of the environmental microbiome. This is because the host microbiome is not affected by MS and EMS, but the environmental microbiome is reduced by EMS.

### Resource Provisioning parameter, *k*

In our previous research [24], we concluded that hosts are unwilling to provide resources (*k* = 0) when the following three conditions are simultaneously met: when the intensity of microbial selection (MS) and parental inheritance (PI) are both low, and parental contribution to the environment (PE) is high. As discussed previously [24], under these conditions, beneficial microbes acquired by a parent that provides resources by sacrificing fitness are not typically inherited by offspring; however, because hosts make a large microbial contribution to the environment, hosts with beneficial microbes release these microbes into the environment so that other hosts are able to acquire these beneficial microorganisms without sacrificing any fitness. So a host that sacrifices its own fitness to promote the presence of good microorganisms will not benefit from this sacrifice because it will not pass on the microbes to its offspring. Instead, by contributing these beneficial microbes to the environment, it will benefit other hosts that have not sacrificed any fitness. Over evolutionary time, *k* will tend to zero.

Here, we show that in the presence of EMS, hosts have to sacrifice more resources (and therefore, fitness) to ensure that they recruit beneficial microbes. This can be seen by comparing the heatmaps in Fig. 4a and Fig. 4b. As can seen, particularly when *S_MS_ >* 1 and *S_HS_ >* 1, mean values of *k* tend to be higher in the presence EMS (i.e., *S_EMS_*= 100) than when EMS is absent (i.e., *S_EMS_* = 1). We also plotted a barplot of the mean *k* under different intensities of EMS (Appendix B: Fig.). We performed an analysis of variance (ANOVA) on *k* at different EMS intensities to confirm the difference in the mean values of *k* (*p <* 0.001).

### Composition of Microbiomes in Environment

When considering the composition of microorganisms in the environment, we have classified microorganisms in two ways. First, we classified them according to their fitness in the environment: when the environmental fitness *>* 0, the environment positively selects that microbe; conversely, when environmental fitness *<* 0, the environment has a negative selective effect on the microbe; and finally, when environmental fitness = 0, the environment is neutral with respect to the fitness of the microbe. Second, we classified them according to their impact on the survival of the host (i.e., when the contribution to the host fitness *>* 0, = 0, or *<* 0).

#### First Classification: Microbial Fitness in the Environment

Let’s look at the first classification (Fig. 5a). The proportion of microorganisms that have positive fitness in the environment increases as the intensity of EMS increases. However, this trend is also affected by the parental contribution. The higher the parental contribution, the lower the proportion of positive microorganisms (e.g., *S_EMS_* = 100 and *S_MS_* = 1000). This is because, in the case of high parental contribution, the host provides the environment with a large number of microorganisms that are suitable for survival in the host, and these microorganisms are not necessarily positively selected for in the environment.

**Figure 5:**
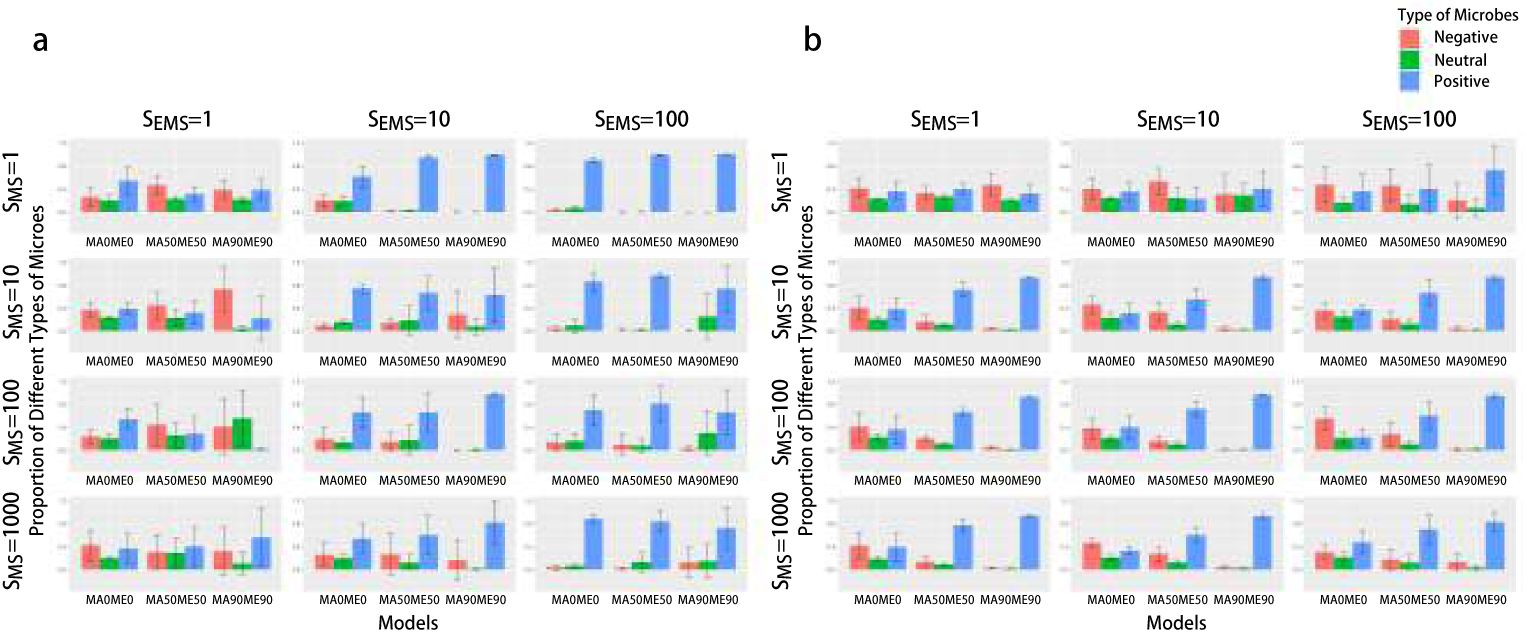
Composition of microbiomes under different models in environment. Each barplot represents its corresponding combination of EMS and MS. There are three levels of EMS: 1, 10, 100, and four levels of MS: 1, 10, 100, 1000. For each barplot, the horizontal axis represents different combinations of parental contribution rates: *MA0ME0*, *MA50*ME50*, *MA90ME90*. *MAnMEm* represents a parental contribution rate of n% to the host and m% to the environment. The vertical axis represents the microbial proportions, and each bar represents the average value of five repetitions. The lines at each bar represent ± one standard deviation. **a** The color coding is as follows: red represents negative microbes (environmental microbial fitness *<* 0), green represents neutral microbes (environmental microbial fitness = 0), and blue represents positive microbes (environmental microbial fitness *>* 0). **b** The color coding is as follows: red represents negative microbes (host fitness *<* 0), green represents neutral microbes (host fitness = 0), and blue represents positive microbes (host fitness *>* 0).

#### Second Classification: Microbial Contribution to Host Fitness

In the second classification (Fig. 5b), we can observe that microorganisms beneficial to the host increase in the environment with the addition of MS. Again, this is influenced by the parental contribution rate. The higher the parental contribution rate, the more microorganisms in the environment that are beneficial to the host. However, surprisingly, EMS does not seem to have much effect.

### Composition of Microbiome in Hosts

When we look at microbes within their hosts, we find that microorganisms which are beneficial to the host in the host increase with increasing parental contribution when there is no EMS (Fig. 6). At the same time, an increase in MS intensity also leads to an increase in positive microorganisms in the host. This is in line with our findings in our previous research. Similarly, differences in EMS intensity have no strong effect on the relative proportions of positive, neutral, or negative microbes.

**Figure 6:**
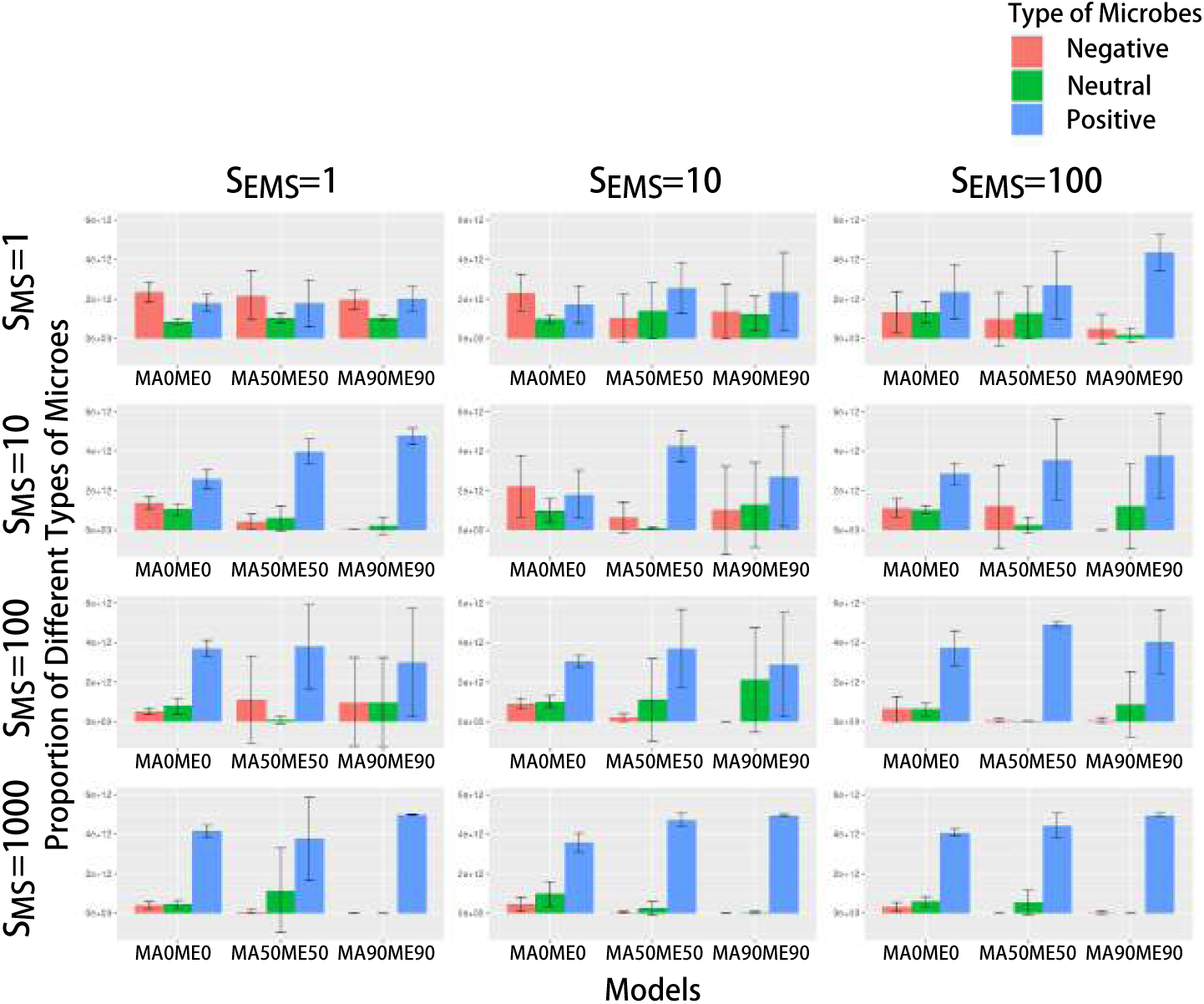
Composition of microbiomes under different models in hosts. This figure is plotted in the same way as Fig. 5. And the color coding is as follows: red represents negative microbes (host fitness *<* 0), green represents neutral microbes (host fitness = 0), and blue represents positive microbes (host fitness *>* 0).

### Fitness of Hosts and Microbiomes

In line with expectations, the change in host fitness increases with increasing parental contribution and HS strength (Fig. 7a), corresponding to the results of our previous research [24]. However, corresponding to the previous, the change in EMS strength does not cause much change.

**Figure 7:**
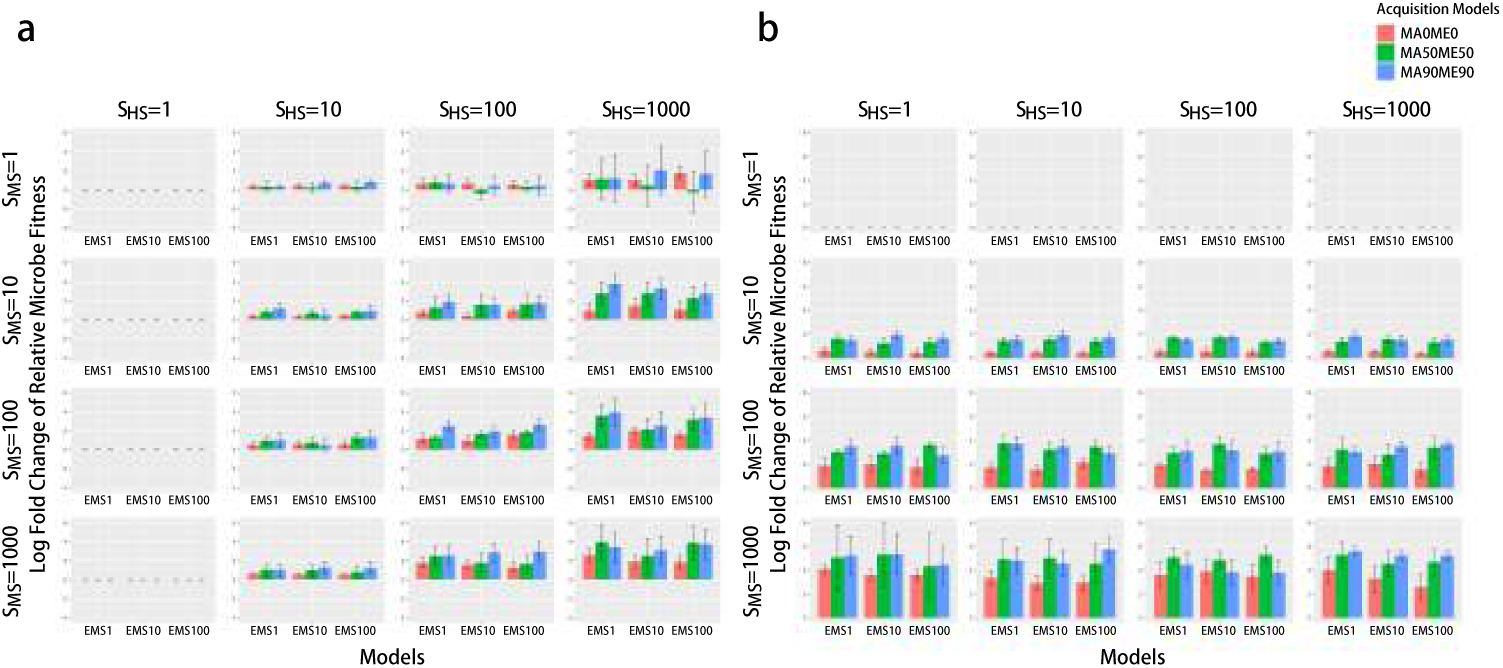
The average log-fold relative change in host fitness at the start and end of the simulations under different intensities of EMS with RPP. Each barplot represents its corresponding combination of HS and MS. There are four levels of HS: 1, 10, 100, 1000, and four levels of MS: 1, 10, 100, 1000. The vertical axis represents the logarithmic change in average host fitness relative to the initial level. The horizontal axis represents three levels of EMS: 1, 10, 100. *MSEn* represents *S_EMS_* = *n*. The lines at each bar represent ± one standard deviation. Different colors represent different combinations of parental contribution rates.

Secondly, and also corresponding to the results of our previous research, the change in microbiome fitness also increases with the increase in the strength of MS (Fig. 7b). However, changes in EMS do not play a role.

## Discussion

In a previous study [24], we extended the Zeng et al. [25] selection model by including the RPP. However, in nature, the environment also has a selective influence on microorganisms, but our previous model was not able to describe this. Therefore, in this study, we included environmental microbial selection to model the impact of the environment on microbial diversity, and host and microbial fitnesses.

First, our results show that *α*-diversity in the environment decreases as the intensity of EMS and host influence on the microbial composition of the environment increases (Fig. 2). This is exactly what we expect to see: EMS causes the frequency of microorganisms with low fitness in the environment to decrease over time. It is true that in extreme environments, microbial diversity usually decreases due to extreme conditions. For example, high salt environments have limited microbial diversity due to a combination of high salt concentration, temperature, *pH*, and other factors [26]; soda lakes have high *pH* due to high levels of sodium carbonate, which leads to low microbial diversity [27]; and the acidification process of a lead/zinc mine while it is being mined leads to lower microbial diversity [28].

Surprisingly, in the presence of both EMS and MS, environmental *α*-diversity can increase. There are microorganisms in the environment that would not otherwise survive in the environment but are able to proliferate in the host. Since the environmental microbial community can be influenced by contributions from the host microbiome in our model, these microbes can also survive in the environment where they may be potentially less fit. In this way, the *α*-diversity in the environment is enhanced. This means that some microorganisms in the environment survive because of the existence of their hosts. If the hosts of such microorganisms goes extinct, the microorganisms are also likely to go extinct, unless alternate hosts/environmental niches are available [29].

The *α*-diversity of the host’s microbiome is influenced by both the environment in which the host lives and the internal environment of the host. Our results show that if the *α*-diversity in the environment decreases, the *α*-diversity of the host’s microbiome also decreases (Fig. 2 and Fig. 3). Moreover, this relationship becomes more apparent when the environment has a significant impact on the microbiome of the host. For example, some researchers have found, through an examination of the microbiomes and genotypes of 1046 healthy individuals, that the environment has a greater influence than host genetics in shaping the human gut microbiota [6]. Numerous studies have shown that microbial diversity is lower in urban areas compared to rural areas [30–32]. Furthermore, investigations on the gut microbiome of healthy adults have revealed that the diversity of gut microbiome is lower in urban populations from the United States and Italy compared to rural populations from Papua New Guinea, Native American communities, Malawi, as well as hunters from Tanzania and the Amazon region [33–35]. Researchers conducted a microbial investigation of the living environments and intestines of fullterm infants in urban and rural areas [36]. And the results revealed that microbial diversity in rural living environments was significantly higher than in urban ones. Additionally, the diversity of intestinal microbiota in rural infants was also higher compared to urban infants [36].

Another interesting result that emerges from our model is the increase in the amount of fitness a host is willing to sacrifice to recruit/retain microbes (measured by *k*) in the presence of EMS. Our model implies that in certain environments, especially under relatively harsh conditions, the evolution of obligate symbiotic relationships is more likely to occur. The host provides resources to the microbes, and in return, the microbes reciprocate by providing benefits to the host. This mutualistic interaction between the host and microbes promotes co-evolution and enhances the fitness of both parties in challenging environments. A notable example of this is seen in tropical corals and their endosymbiotic dinoflagellate algae, where the low nutrient characteristics of open tropical oceans result in low primary productivity [37]. However, corals obtain photosynthetically fixed carbon from algae, while the algae acquire the necessary inorganic compounds (nitrogen) from the corals [38, 39]. Importantly, this symbiotic relationship cannot be sustained solely through heterotrophic feeding and nutrient uptake from seawater [40, 41]. *Rhizobium sullae* and *Hedysarum* plants also have a mutually beneficial symbiotic relationship. *Rhizobium sullae* is not able to survive the extreme drought conditions that *Hedysarum* plants can tolerate [42], but it can survive the symbiotic relationship by providing nitrogen to *Hedysarum* plants for nitrogen fixation [43–45]. This point also validates what was mentioned earlier, that some microorganisms survive because of their hosts. This is an interesting hypothesis to test: do we expect to see more host resource provisioning in harsher environments?

Based on the composition of microorganisms in the environment, we know that an increase in the intensity of EMS in the environment leads to the survival of microorganisms that can adapt to the characteristics of the environment. This phenomenon is particularly evident in extreme environments. For instance, researchers sampled 925 geothermal springs (pH *<* 1–9.7 and 13.9–100.6 °C) in The Taupō Volcanic Zone of the North Island of New Zealand [46]. Through 16S rRNA gene amplicon sequencing, they found that Proteobacteria and *Aquificae* dominate the microbial communities. Among the detected proteobacterial genera, the most abundant ones are mainly aerobic chemolithoautotrophs that utilize sulfur species and/or hydrogen for metabolism. *Acidithiobacillus*, in particular, is the most abundant and widespread genus. *Acidithiobacillus* is known as a mesophilic and acidophilic autotroph that utilizes reduced sulfur compounds, iron, and/or hydrogen as its growth energy sources [47]. Similarly, terrestrial *Aquificae* are primarily microaerophilic chemolithoautotrophs that oxidize hydrogen or reduced sulfur compounds [48].

Furthermore, in our results, EMS does not significantly influence the abundance of beneficial microbes for the host in the environment. This indicates that the survival of host-beneficial microbes in the environment is primarily controlled by their respective hosts rather than the environment itself. This aligns with our previous hypothesis [24] that the survival of certain microbes is contingent upon their association with specific hosts. Therefore, in order to conserve microbial diversity, the conservation efforts for their hosts should not be overlooked [49].

Finally, in our simulations, EMS does not influence the composition of the host microbiome, host fitness, or microbe fitness within the host. There are two possible reasons for this outcome. Firstly, RPP (resource provisioning process) plays a significant and compensatory role within the host. As concluded in the previous study [24], RPP allows the maintenance of the proportion of beneficial microorganisms in the host and enhances host fitness. At the same time, microbial fitness in the host is also enhanced by MS. Secondly, the current model assumes EMS is neutral for the host, which means EMS doesn’t deliberately increase the proportion of positive or negative OTUs (for hosts) in the environment. To validate these findings, future computational models can explore the effects on the composition of the host microbiome, host fitness, and microbe fitness within the host by introducing specific EMS scenarios, such as creating environments that are 100% beneficial or harmful to the host.

## Materials and Methods

The details of neutral and selection models have been described in previous study [25, 50].

### EMS

In the EMS model, the contribution to environmental microbial fitness of OTUs can be calculated by:

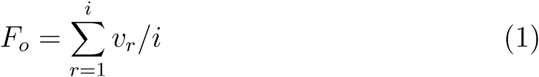

where *F_o_*represents the contribution to environmental microbial fitness of OTU *o*, *v_r_* represents the value of the *r* th trait of OTU *o* which can be -1, 0, and 1, *i* represents the number of traits of OTU *o*.

The probability that an OTU is acquired by the host can be calculated by:

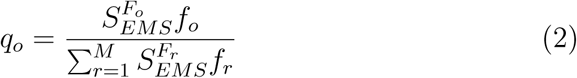

where *q_o_*represents the probability of OTU *o*, *F_o_*represents the environmental microbial fitness of OTU *o*, *f_o_* represents the frequency of OTU *o* in the mixed environment, *M* represents the number of OTUs in the environment, *S_EMS_* represents the intensity of EMS.

### Calculation of Microbial Diversity

In this study, *α*-diversity and *γ*-diversity (the overall microbial diversity from all hosts in the population) are calculated by scaled Shannon-Wiener index [51]:

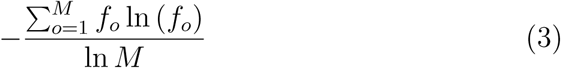

where *M* represents the total number of OTUs in the environment, *f_o_* represents the frequency of OTU *o* in microbial community.

And *β*-diversity (the change in microbial diversity between different hosts) is calculated by Bray-Curtis index [52]:

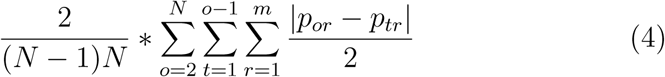

where *N* represents the number of hosts, *m* represents the total number of OTUs in all hosts, *p_or_* and *p_tr_* represent the frequency of OTU *r* in Host *o* and Host *t*.

### Model Implementation

In this paper, we incorporated EMS into the model discussed in our previous research. Before simulating the model, we assigned an additional random environmental fitness to each trait to calculate the fitness of microbes in the environment. We used the same parameters as in our previous research and combined different intensities of EMS. All graphs were plotted using R language, while all simulations were performed using Java language. The model was run on the supercomputer system NeSI. The model was run on the supercomputer system NeSI.

## Supporting information

The simulation results without RPP and other results with different HS intensities.

## Acknowledgements

We thank Qinglong Zeng for assistance with the original code, and Steve Wu and Anna Santure for helpful comments on an earlier draft of this work. Funds for this research were provided by The University of Auckland, New Zealand.

## Data Availability Statement

The code and dataset of this study are available at github.com/zjzxiaohei/Environmental-Selection-Model.git.

