## Supplementary material for "Models of microbiome evolution incorporating environmental microbial selection": The simulation results without RPP and other results with different HS intensities.

### Supplemental files 1: Figure S1

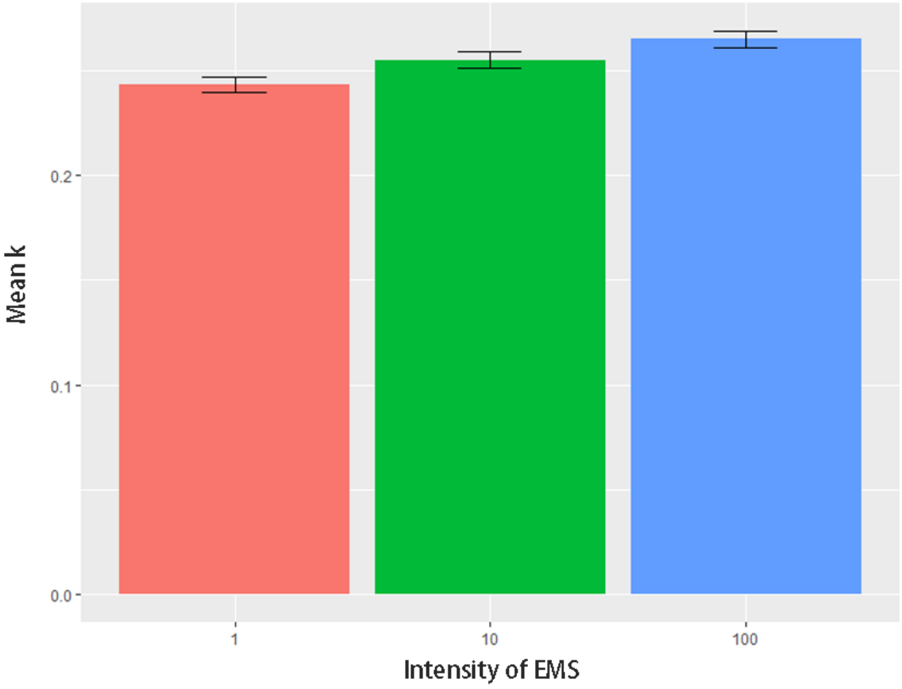


Figure S1: The mean $k$ under different intensity of EMS. The vertical axis represents the mean $k$. The horizontal axis represents four levels of EMS: 1, 10, 100. The mean $k$ is calculated from the data obtained when both HS and MS are present ($S_{HS} \neq 0$ and $S_{MS} \neq 0$). The lines at each bar represent $\pm$ one standard error.

### Supplemental files 2: Figure S2

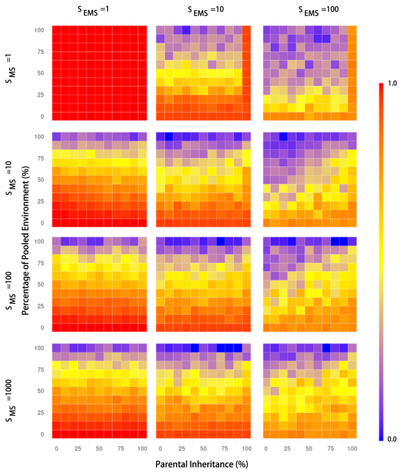


(a) $S_{HS} = 1$

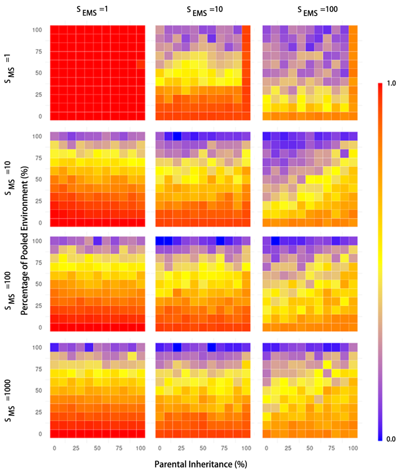


(b) $S_{HS} = 10$

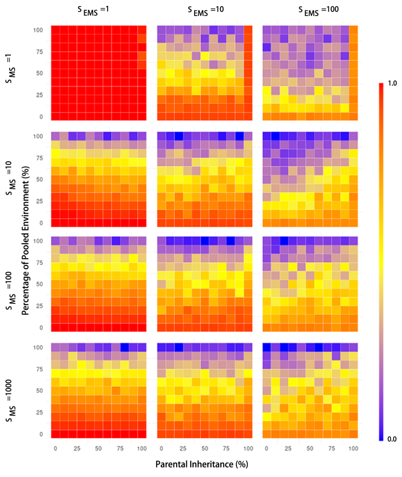


(c) $S_{HS} = 100$

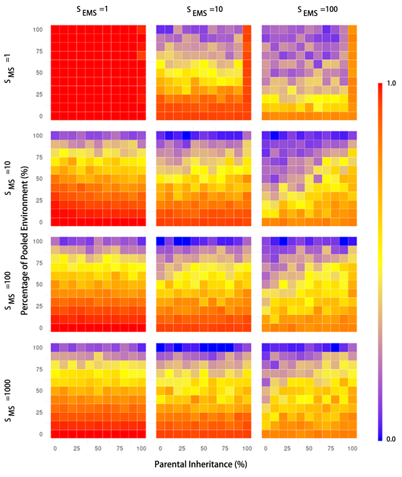


(d) $S_{HS} = 1000$

Figure S2: Pattern of $\alpha$-diversity in the environment without RPP.

### Supplemental files 3: Figure S3

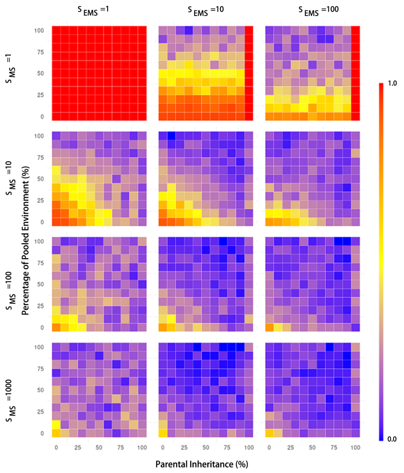


(a) $S_{HS} = 1$

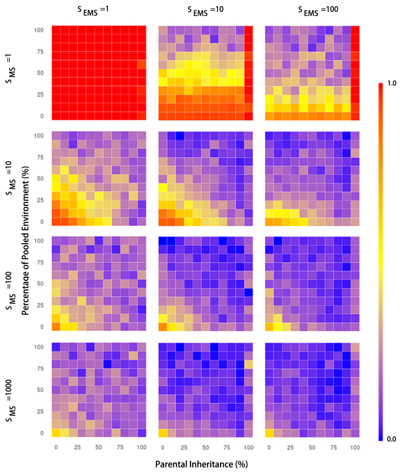


(b) $S_{HS} = 10$

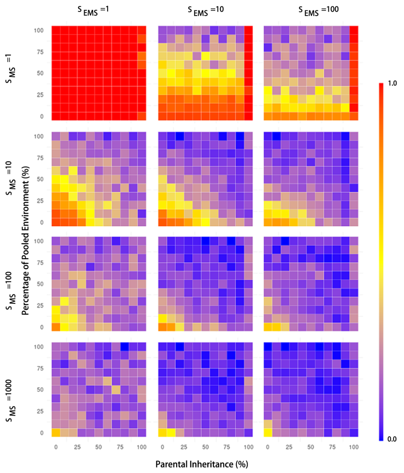


(c) $S_{HS} = 100$

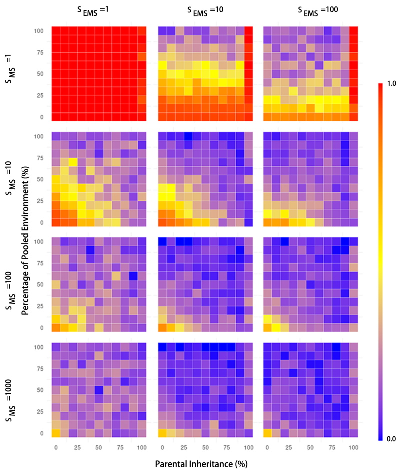


(d) $S_{HS} = 1000$

Figure S3: Pattern of $\alpha$-diversity in the hosts without RPP.

### Supplemental files 4: Figure S4

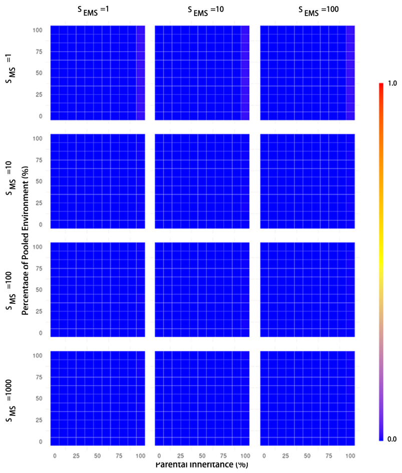


(a) $S_{HS} = 1$

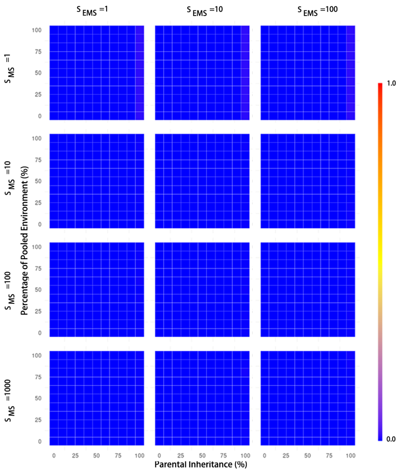


(b) $S_{HS} = 10$

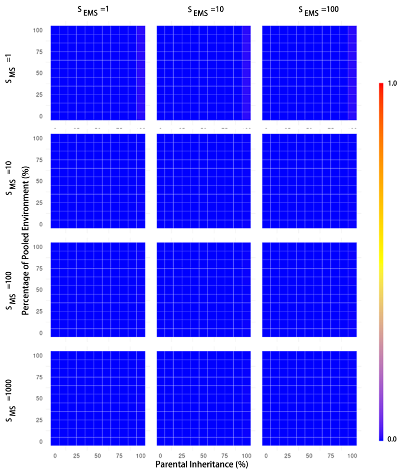


(c) $S_{HS} = 100$

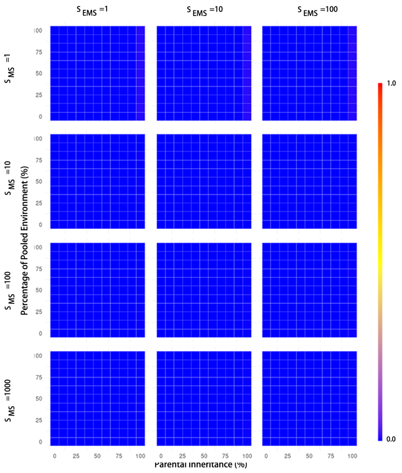


(d) $S_{HS} = 1000$

Figure S4: Pattern of $\beta$-diversity in the hosts without RPP.

### Supplemental files 5: Figure S5

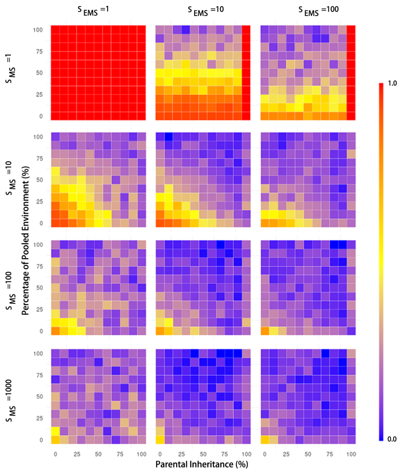


(a) $S_{HS} = 1$

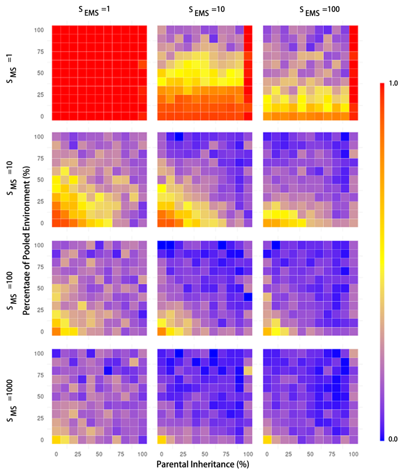


(b) $S_{HS} = 10$

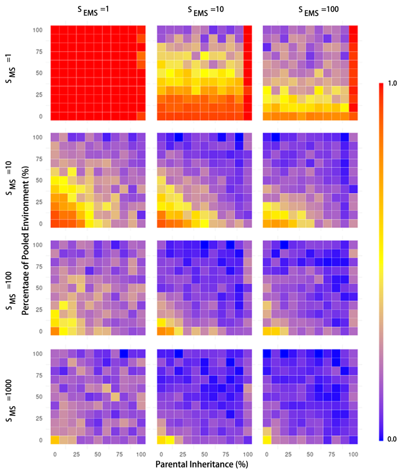


(c) $S_{HS} = 100$

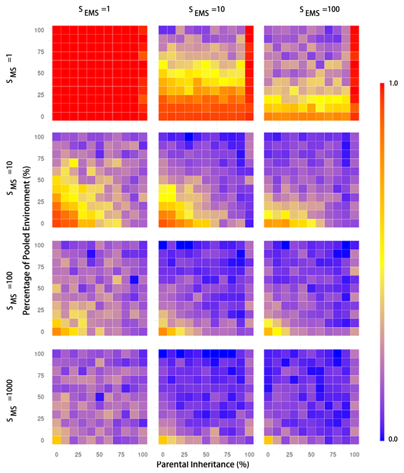


(d) $S_{HS} = 1000$

Figure S5: Pattern of $\gamma$-diversity in the hosts without RPP.

### Supplemental files 6: Figure S6

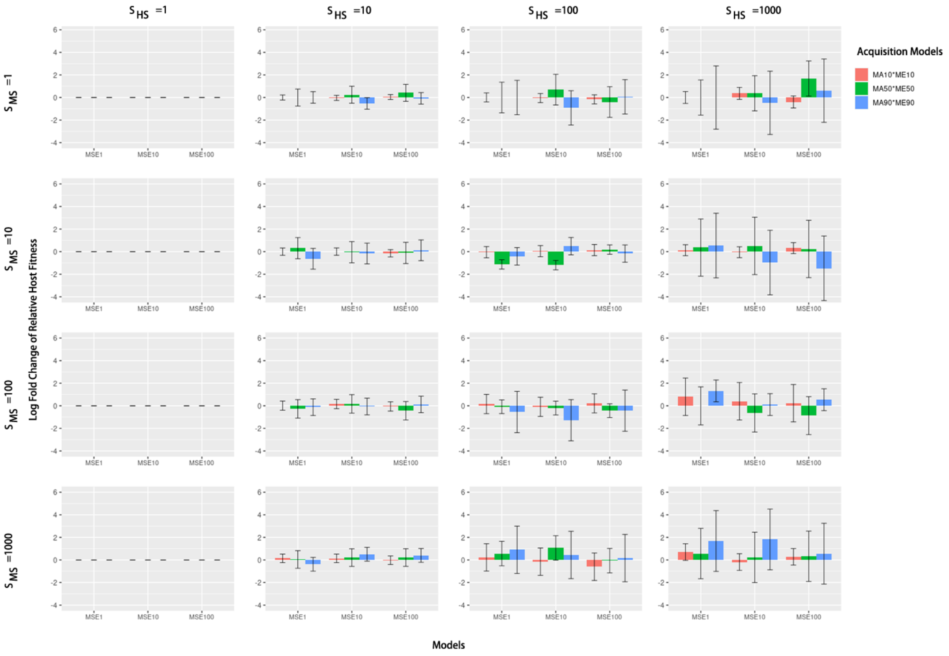


Figure S6: The variation of host fitness under different models without RPP.

### Supplemental files 7: Figure S7

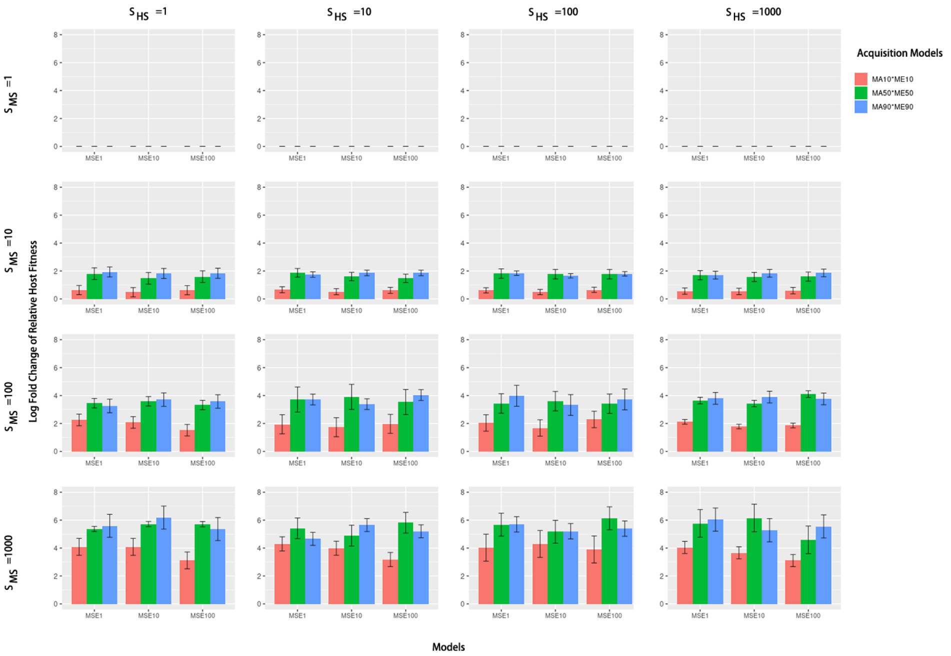


Figure S7: The variation of microbiome fitness under different models without RPP.

### Supplemental files 8: Figure S8

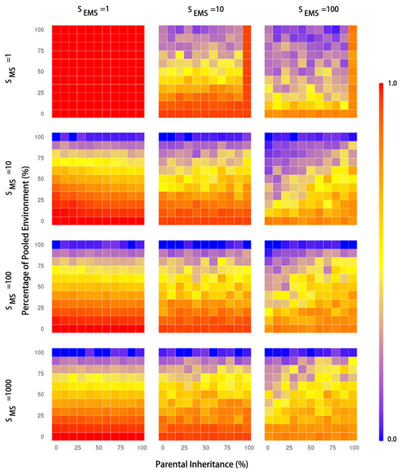


(a) $S_{HS} = 1$

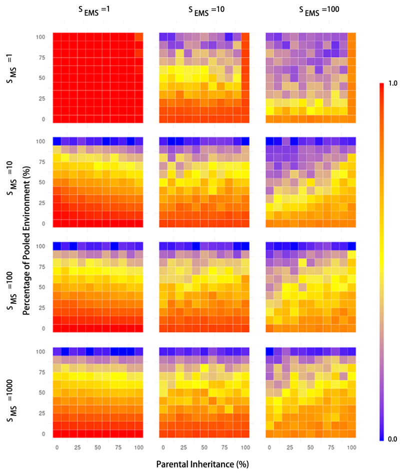


(b) $S_{HS} = 10$

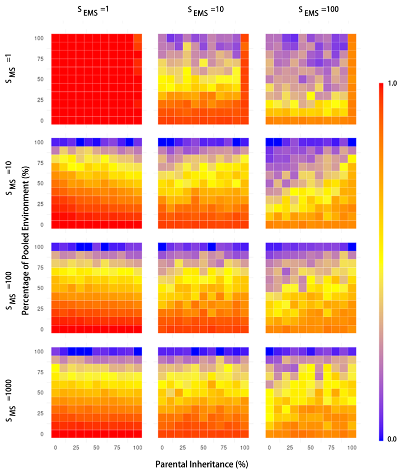


(c) $S_{HS} = 100$

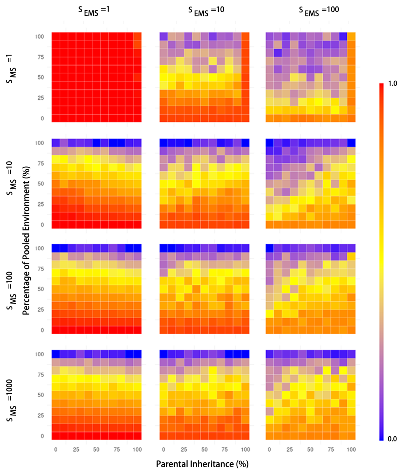


(d) $S_{HS} = 1000$

Figure S8: Pattern of $\alpha$-diversity in the environment with RPP.

### Supplemental files 9: Figure S9

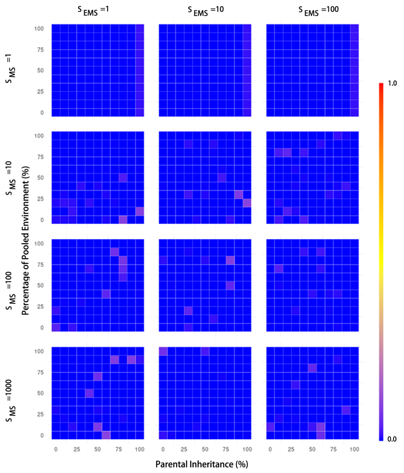


(a) $S_{HS} = 1$

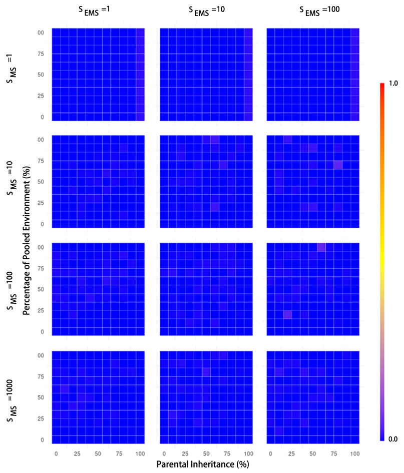


(b) $S_{HS} = 10$

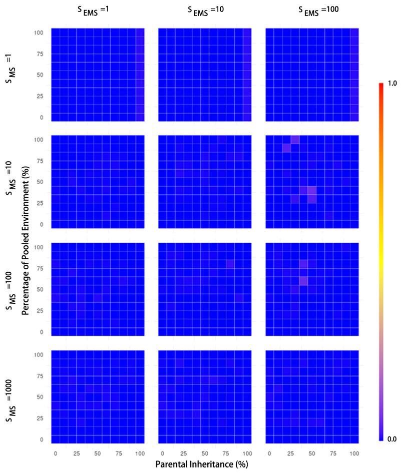


(c) $S_{HS} = 100$

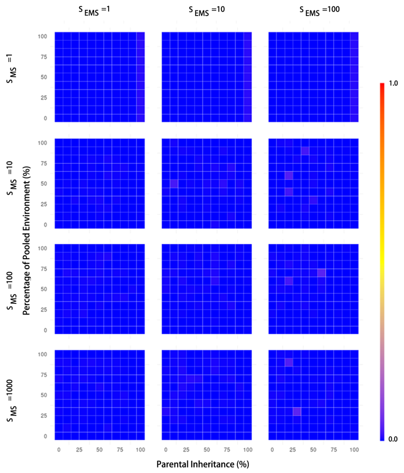


(d) $S_{HS} = 1000$

Figure S9: Pattern of $\beta$-diversity in the hosts with RPP.

### Supplemental files 10: Figure S10

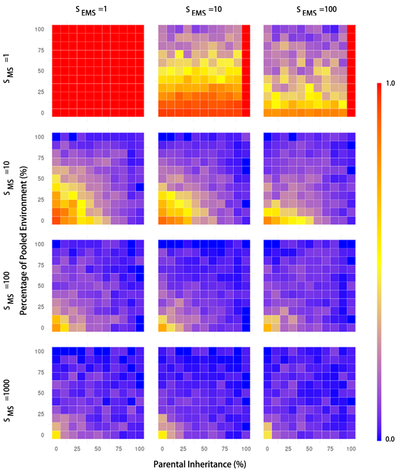


(a) $S_{HS} = 1$

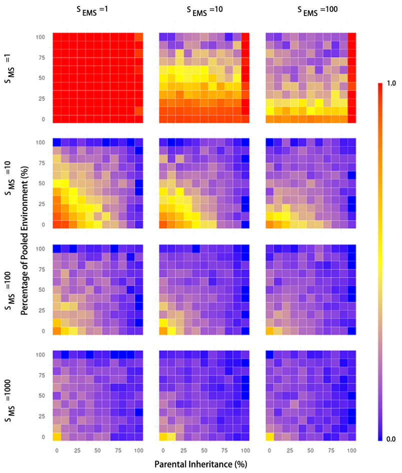


(b) $S_{HS} = 10$

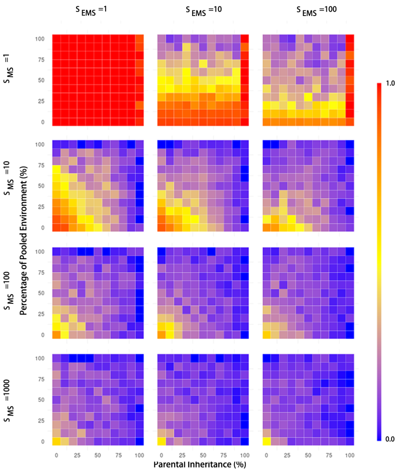


(c) $S_{HS} = 100$

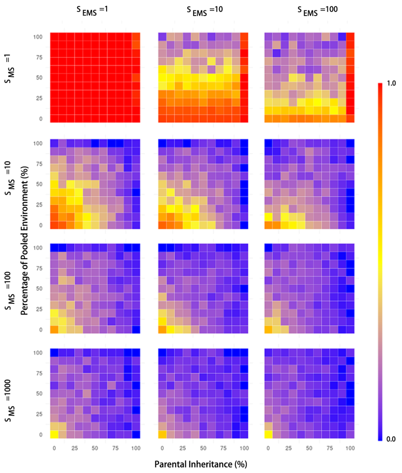


(d) $S_{HS} = 1000$

Figure S10: Pattern of $\alpha$-diversity in the hosts with RPP.

### Supplemental files 11: Figure S11

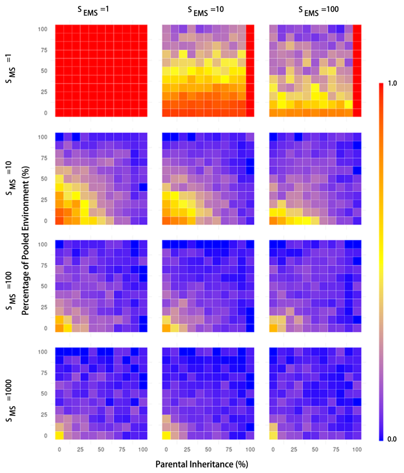


(a) $S_{HS} = 1$

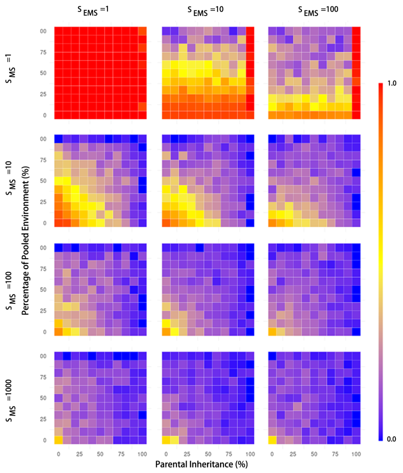


(b) $S_{HS} = 10$

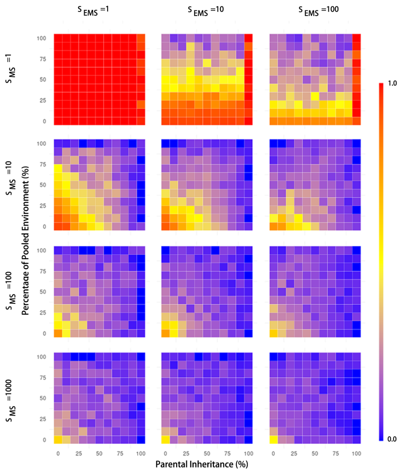


(c) $S_{HS} = 100$

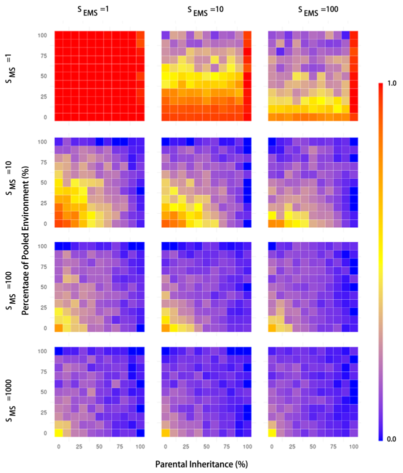


(d) $S_{HS} = 1000$

Figure S11: Pattern of $\gamma$-diversity in the hosts with RPP.

### Supplemental files 12: Figure S12

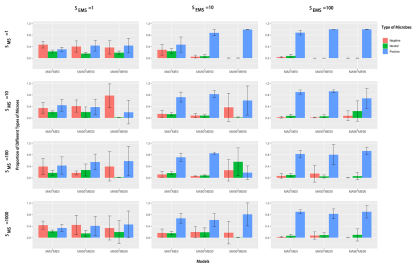

(a) $S_{HS} = 1$

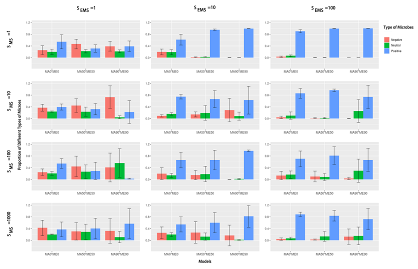

(b) $S_{HS} = 10$

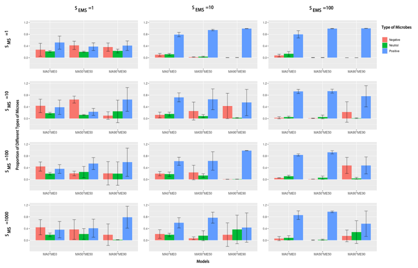

(c) $S_{HS} = 100$

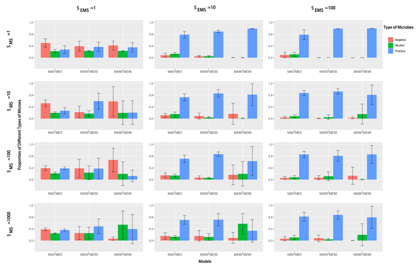

(d) $S_{HS} = 1000$

Figure S12: Composition of microbiomes under different models in environment. The color coding is as follows: red represents negative microbes (environmental microbial fitness $< 0$), green represents neutral microbes (environmental microbial fitness $= 0$), and blue represents positive microbes (environmental microbial fitness $> 0$). The lines at each bar represent $\pm$ one standard deviation.

### Supplemental files 13: Figure S13

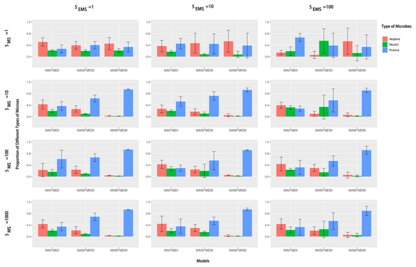

(a) $S_{HS} = 1$

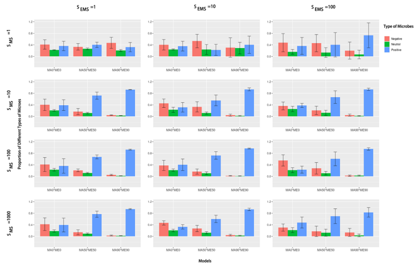

(b) $S_{HS} = 10$

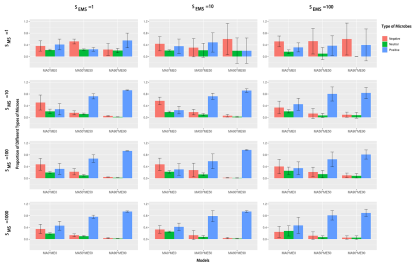

(c) $S_{HS} = 100$

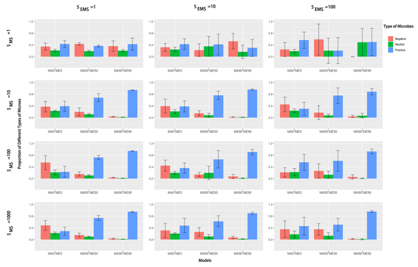

(d) $S_{HS} = 1000$

Figure S13: Composition of microbiomes under different models in environment. The color coding is as follows: red represents negative microbes (host fitness $< 0$), green represents neutral microbes (host fitness $= 0$), and blue represents positive microbes (host fitness $> 0$).

14 # Supplemental files 14: Figure S14

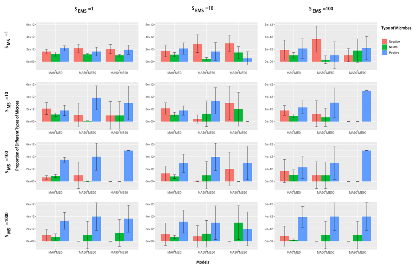

(a) $S_{HS} = 1$

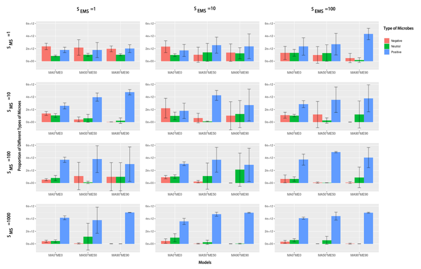

(b) $S_{HS} = 10$

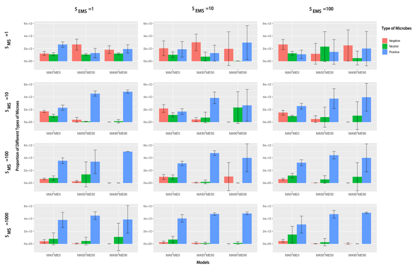

(c) $S_{HS} = 100$

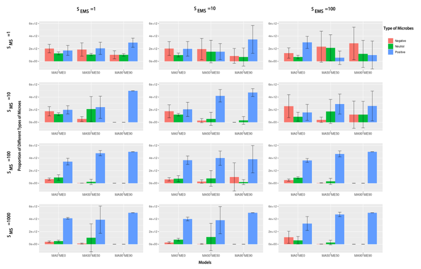

(d) $S_{HS} = 1000$

Figure S14: Composition of microbiomes under different models in hosts. The color coding is as follows: red represents negative microbes (host fitness $< 0$), green represents neutral microbes (host fitness $= 0$), and blue represents positive microbes (host fitness $> 0$).
